# Long-term memory formation impacts dietary energy intake but not metabolic rate in honeybees

**DOI:** 10.1101/2025.02.17.638520

**Authors:** Cecylia M. Watrobska, Jonathan Codd, Steven J. Portugal, Ellouise Leadbeater

## Abstract

The brain is energetically expensive. Energy availability may, therefore, determine whether costly cognitive processes such as long-term memory can be expressed. However, there is a limited understanding of the metabolic costs associated with long-term memory formation. Here, we explored the potential induced costs of long-term memory formation using honeybees (*Apis mellifera*) as a model species. We monitored the sucrose intake of bees over the 20-hour period following a classical spaced olfactory conditioning protocol that induced long-term memory formation, relative to a control group that experienced the same reward schedule but no odour pairing. Bees in the experimental treatment drank significantly more sucrose than controls. We then tested whether the increased energy demands of long-term memory formation showed parallel increases in metabolic rate, by measuring carbon dioxide production in groups of bees at four timepoints following conditioning (1-hour, 4-hours, 24-hours and 72-hours). We found no change in metabolic rate between learning and control groups across all time points. Our results suggest that the cost of long-term memory formation does not translate to increases in metabolic rate, despite an increase in dietary energy intake. Constraints on the expression of cognitive traits may, therefore, be unlikely when energy is unrestricted and can be compensated.

## Introduction

Neural tissue is fundamental to the expression of cognitive traits, yet its maintenance and use are energetically expensive (Attwell & Laughlin, 2001; Niven & Laughlin, 2008). Accordingly, studies across taxa have documented negative correlations between brain size and other costly organs or traits (e.g., gut size or immunity; Aiello & Wheeler, 1995; Isler & van Schaik, 2006; Kotrschal et al., 2013), suggesting that energy is diverted to the maintenance of a relatively larger brain. The highest energy investment is required for synapse maintenance and synaptic transmission (Harris, Jolivet & Attwell, 2012). Approximate estimates based on mammalian grey matter, a neuronally-dense area of the brain, suggest that almost half of the brain energy budget is invested in action potentials that process and transmit information (Attwell & Laughlin, 2001; Harris, Jolivet & Attwell, 2012). Predictions suggest, therefore, that energy demands may further increase when cognitive traits are expressed (Burns, Foucaud & Mery, 2010; Dukas, 1999).

Long-term memory (LTM) is a physiologically distinct memory phase that persists over multiple days and induces *de novo* protein synthesis and synaptic re-organisation in the brain (Davis, 2011; Menzel, 2012; Tully et al., 1994). In honeybees (*Apis mellifera*), LTM formation can be induced following spaced trials of associative conditioning, in which a novel stimulus is paired with a reward and presented over successive trials at ∼10-minute intervals (Menzel, 2001; Menzel et al., 2001). Following spaced conditioning, bees can recall learnt stimuli for approximately 72-hours (Menzel et al., 2001) and show corresponding changes in synaptic complex (microglomeruli) densities in the mushroom bodies, which are integrative centres of the insect brain closely associated with learning and memory (Heisenberg, 1998). For example, honeybees forming olfactory LTM had a relatively higher density of synaptic complexes in the lip region of the mushroom bodies (which receives olfactory input) compared with control bees (Hourcade et al., 2010). Increases in synaptic complex densities have further been identified in other social insects and across learning contexts (visual LTM in bumblebees *Bombus terrestris*, Li et al., 2017; long-term avoidance memory in ants *Acromyrmex ambiguus*, Falibene, Roces & Rössler, 2015).

Changes in synaptic complex densities in response to learning and LTM formation may carry an energetic cost above requirements for baseline maintenance of the brain, because energy is necessary to maintain ion flux, activate proteins at the synapse (e.g., through phosphorylation), and for protein synthesis (Harris, Jolivet & Attwell, 2012; Karbowski, 2019). In the hippocampus of rats (*Rattus norvegicus domestica*), LTM formation requires the release of glycogen from energy reserves, and LTM is impaired when glycogenesis is blocked (Suzuki et al., 2011). More recently, evidence from fruit flies (*Drosophila melanogaster*) suggests that LTM formation requires higher rates of pyruvate consumption in mushroom body neurons, which is a substrate for ATP synthesis (Plaçais et al., 2017). Consistent with this, Plaçais et al. (2017) observed that fruit flies doubled their sucrose intake in the first four hours following an aversive olfactory spaced conditioning paradigm compared with flies in a control group, suggesting that cellular energy demands of LTM formation are compensated by increasing dietary energy intake. Consumption in flies that were exposed to a massed conditioning protocol (which uses short intervals between trials to induce anaesthesia resistant memory that is not reliant on protein synthesis, Tully et al., 1994) did not differ from flies in an untrained (control) group. Thus, the increase in consumption appeared to be driven by physiological changes accompanying LTM formation. Overall, these studies suggest LTM formation places increased energy demands on an individual.

In keeping with this, evidence suggests that investing in LTM formation may lead to trade-offs with other traits. For example, exposure to a spaced olfactory conditioning protocol correlated with reduced tolerance to desiccation in fruit flies (Mery & Kawecki, 2005) and reduced survival in honeybees (Jaumann, Scudelari & Naug, 2013). Conversely, restricting energy availability may impair LTM formation. In fruit flies, starvation for 21-hours before and then 24-hours after aversive spaced conditioning resulted in flies failing to form LTM compared with satiated control groups (Plaçais & Preat, 2013), and juvenile nematode mutants with restricted dietary intake (*Caenorhabditis elegans* mutant *eat-2*) were also unable to form LTM (Kauffman et al., 2010).

In the present study, we asked whether there is an induced, measurable effect on demand for dietary energy intake associated with LTM formation in honeybees (*A. mellifera*), and whether this cost is reflected in changes in metabolic rate. We used honeybees as a model because memory is thought to be closely related to foraging efficiency in this species, whereby foragers vist thousands of flowers each day and must form long-term memories of rewarding patches, flower species and foraging routes (Menzel et al. 2001). Honeybees also have relatively larger mushroom bodies (integrative regions of the brain associated with learning and memory) compared with fruit flies (Menzel, 2012) and LTM has been well-established and can be recalled 72 hours post-conditioning (Menzel et al., 2001). We first explored whether individuals increase their sucrose consumption in the 20-hours following a spaced olfactory conditioning protocol, by measuring consumption in honeybees at 10-minute intervals. We then investigated whether LTM formation resulted in changes to carbon dioxide production in groups of bees, as a proxy for standard metabolic rate. Metabolism maintains energy homeostasis and responds to variation in energy demands (Brown et al., 2004). Measuring metabolic rate therefore provides a comprehensive tool to measure energy expenditure. Previous work has explored the relationship between memory and resting metabolic rate in chickens (*Gallus gallus domesticus*, Watrobska et al., 2023), but to our knowledge, metabolic rate has not been studied in the context of potential changes associated with energy investment in memory formation.

## Methods

### Overview

We conducted two experiments in which honeybees (*Apis mellifera*) were trained in an olfactory conditioning assay (Experiment 1: n = 82 learners, n = 79 controls; Experiment 2: n = 115 learners, n = 105 unpaired controls, n = 105 full controls) prior to measuring sucrose consumption (Experiment 1) and metabolic rate (Experiment 2).

### Experimental animals

Returning honeybee foragers were collected from hives at the apiary at Royal Holloway, University of London, UK (latitude = 51.423283, longitude = −0.566432; Experiment 1: n = 4 hives August-October 2021; Experiment 2: n = 6 hives, April-May 2023). For Experiment 1 (effects of learning and memory formation on sucrose consumption), bees were collected each morning, taken to the laboratory, placed on ice until immobile (approximately five minutes, Matsumoto et al., 2012), and harnessed into adapted Eppendorf tubes secured with a strip of electrical tape behind the head. Following harnessing, all bees were fed with 1 µL 40% w/w sucrose solution (granulated sugar dissolved in distilled water) and left to acclimatise for one hour at 25°C and 60% RH. For Experiment 2 (effects of learning and memory formation on metabolic rate), bees were collected the night before learning trials (between 16:00-18:00) to account for different energy states of foraging bees before metabolic rate measurements, harnessed as described above and fed to satiation with 30% w/w sucrose solution. We adapted the harnesses by drilling holes in the sides to allow for efficient gas exchange during metabolic rate measurements. The following morning, we fed bees with 1 µL 30% sucrose solution and waited for a further one hour prior to starting experiments.

Before learning trials began, we presented each bee with 50% w/w sucrose solution by touching the antennae (maximum 15 seconds for each antenna), and any bees that did not extend their proboscis (indicating that they were not motivated to participate in trials) were excluded from the experiment (Bitterman et al., 1983). Bees were assigned randomly to treatment groups (Experiment 1: learning or control – full; Experiment 2: learning, control – unpaired odour and reward presentation, or control – full). Following conditioning and sucrose consumption or metabolic rate measurements, we determined the dry body mass of individuals after euthanasia by drying them at 70°C for 48 hours (Hotbox Oven, Gallenkamp, Cambridge, UK).

### Olfactory conditioning protocol

We conditioned bees to associate an odour (Experiment 1: orange/ginger, Experiment 2: orange/ginger/lemon/aniseed 100% pure essential oils, 3 µL, Calmer Solutions) with a sucrose reward using the proboscis extension response assay (Fig. 1A; Bitterman et al., 1983; Menzel et al., 2001). Individual bees were placed into the experimental set-up, which consisted of a clear Perspex arena (25×25×25 cm) from which air was continuously extracted. Odour delivery was controlled using an Arduino-type microcontroller (Orangepip® Kona328) and three solenoid valves, each attached to a 1 mL glass syringe that contained filter paper (1 cm^2^) soaked with the relevant odour (or blank filter paper for unscented air). Solenoid valves received input from an aquarium pump (JBL ProSilent A400 Air Pump).

**Figure 1.**
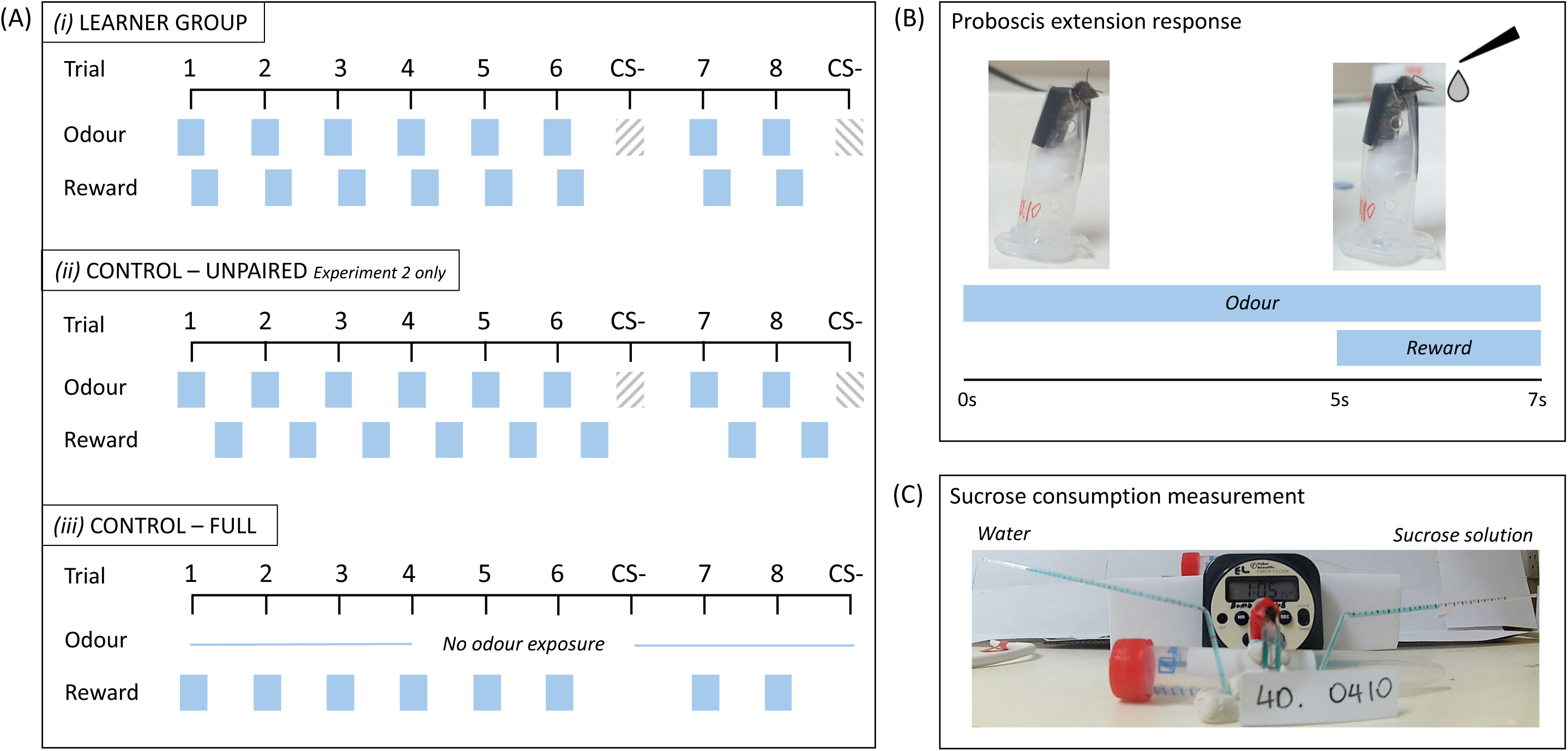
(A) Associative conditioning trials in which honeybees were exposed to an odour paired with a sucrose reward *(i)*, and control groups in which the odour and reward were unpaired to preclude learning *(ii)*, or bees were not presented with the odour but were fed an equivalent volume of sucrose to control for energetic state *(iii)*. In the learner group *(i)*, bees were presented with eight trials of the rewarded odour, and two trials of a different, unrewarded odour (labelled CS-, shaded grey boxes). The inter-trial interval was 10 minutes. In the unpaired control group *(ii)*, odour presentation was spaced by 10 minutes, and sucrose was presented 5 minutes after odour presentation had ended. In the full control group *(iii)*, bees did not enter the experimental setup, and were fed an equivalent reward volume every ten minutes, corresponding to feeding patterns of bees in the learner group. For Experiment 1 we included learner and full control groups, and for Experiment 2 we included learner, unpaired control and full control groups. (B) During a single trial, bees in the learner group were presented with the odour for 7 seconds, and the final 2 seconds of odour presentation overlapped with presentation of the sucrose reward, eliciting extension of the proboscis. (C) Experimental set-up for measuring sucrose consumption in honeybees. Bees remained harnessed in a modified Eppendorf tube secured with a piece of electrical tape following olfactory conditioning. Harnesses were secured above two capillary tubes, giving each bee access to two tubes, one containing 40% (w/w) sucrose solution and the other containing distilled water. Capillary tubes were marked externally every 2.5 µL, and bees were filmed over a 20-hour period on a time-lapse setting to measure consumption. Blue food colouring was added to the solutions to aid visualisation.

Each trial began with 30 seconds of unscented air to acclimatise the subject to the setup, followed by presentation with an odour (conditioned stimulus, CS+) for seven seconds (Fig. 1B). The antennae were stimulated with 50% w/w sucrose solution (unconditioned stimulus, US) for the final two seconds of CS+ presentation, to elicit extension of the proboscis, followed by the bee receiving 0.4 µL 50% sucrose solution as a reward. We considered a positive (conditioned) response to the CS+ to have occurred when the bee extended its proboscis prior to US presentation (Fig. 1B). Each bee underwent eight CS+ trials with an inter-trial interval of ten minutes. We included two additional trials of a second, unrewarded, odour (CS-, between trials 6 and 7 and after trial 8), to check that conditioning was specific to the CS+ (Fig. 1A).

Odours used for the experiments mimic plants that are unlikely to be encountered by foraging bees in SE England, but nonetheless, to account for potential previous associations concerning the odours, in Experiment 1 we discounted any bees that responded positively to the CS+ on the first trial, prior to US presentation (n = 12). In Experiment 2, we presented bees with a single trial of the CS+ odour prior to metabolic rate and conditioning trials, and discounted bees that responded positively. By excluding these bees, we prevented loss of bees from the experiment once initial metabolic rate had been measured. For both experiments, we included a full control group in which bees were not exposed to the conditioning setup or odours but received an equivalent volume of sucrose solution at the same time intervals as bees in the learning treatment. In Experiment 2, we included an additional control group for exposure to the odour, in which bees underwent conditioning trials as described above, but the odour and reward presentations were not paired, such that bees could not form an association between them. Bees in this group were fed an equivalent volume of sucrose solution outside the setup, five minutes after odour presentation. Following conditioning trials, bees remained in their harnesses. For any bees kept overnight or over multiple days (e.g., for memory trials, see below), we fed individuals to satiation with 30% w/w sucrose solution each morning and evening, and kept them in the dark at 25°C and 60% RH (Williams et al., 2013).

### Memory trials

To verify that our classical conditioning protocol successfully induced memory formation, responses to the CS+ were retested 24- or 72-hours (Experiment 1), or 4-, 24- or 72-hours (Experiment 2) after conditioning trials. Individual bees participated in memory trials at only one timepoint. We first checked sucrose responsiveness with 50% w/w sucrose solution, and only bees that responded positively proceeded to memory trials (n = 9 and n = 12 bees excluded for Experiments 1 and 2, respectively). For memory trials, bees were presented with the CS+ and CS-odours in a randomised order, both unrewarded, and we recorded extension of the proboscis. Bees in both control groups were included in memory trials, to check that responses in the learners were above those expected at random in the control groups.

### Experiment 1: Sucrose consumption measurement

We modified the capillary feeding (CAFÉ) assay to measure the volume of sucrose solution consumed by individual bees following olfactory conditioning trials. The CAFÉ assay uses microcapillary tubes to measure precise volume changes and was originally developed for use with fruit flies but has recently been adapted for honeybees (Ja et al., 2007; Reade, Katz & Naug, 2016). Microcapillary tubes (World Precision Instruments, internal diameter: 1.12mm, length: 152mm) were bent into shape using a butane torch (‘U-bend’ at 21mm from the feeding end of the tube, and a further 120° bend at 18mm above the ‘U-bend’; Fig. 1C; Reade, Katz & Naug, 2016) and marked externally every 2.5 µL. Tubes were filled with ∼130 µL of 40% w/w sucrose solution or distilled water dyed with blue food colouring (Navy Blue Liquid Colour, Rainbow Dust Colours, UK, 1 μL/mL) to aid visualisation, and sealed with a drop of mineral oil at the non-feeding end to minimise evaporation.

Consumption trials began immediately after learning trials finished and lasted for 20 hours. Individual bees remained in their modified Eppendorf harnesses and were secured horizontally above two microcapillary tubes containing sucrose and water, such that they could access both tubes simultaneously (Fig. 1C). All trials were filmed on a time-lapse setting (Akaso EK7000 Pro 4K Action Camera, filmed at one frame per minute), and we watched the videos and recorded the volume of sucrose and water consumed every ten minutes to the nearest 2.5 µL, with the observer blinded to treatment. Observations were stopped prematurely if an individual *(i)* died during the trial, *(ii)* was no longer able to reach the microcapillary tubes (e.g., by crawling inside the Eppendorf harness), or *(iii)* had depleted all sucrose or water from the microcapillary tube before the end of the trial (Fig. S1). Videos were analysed by a single observer to ensure consistency.

### Experiment 2: Metabolic rate measurement

We pooled bees into groups of five individuals and measured the rate of carbon dioxide production (*V̇*CO_2_) as a proxy for standard metabolic rate (Lighton, 2008). Each group was first measured immediately before the conditioning protocol, to determine a baseline standard metabolic rate measurement, and then at 1-, 4-, 24- or 72-hours after conditioning, but prior to memory trials. Each group consisted of the same individuals throughout the experiment.

Our open-flow respirometry setup pulled CO_2_-free air (soda lime, Intersorb Plus, Intersurgical) into the chamber (140 mL opaque plastic box covered and kept in darkness to minimise stress and movement, dimensions: 8.5×8.5×5.3 cm) at approximately 225 mL min^-1^, before being scrubbed of water vapour (anhydrous indicating Drierite, W. A. Hammond Drierite Co. Ltd., Ohio, USA) and entering the CO_2_ analyser (Foxbox Field Gas Analysis System, Sable Systems, Las Vegas, USA). Measurements lasted approximately 10 minutes (mean±S.E.M. = 9.97±0.05 minutes), allowing enough time for all air in the chamber to be fully replaced once and the trace to settle. Bees remained in their harnesses for the duration of the metabolic recording to minimise any effects of movement. All experiments were conducted at 25°C and 60% RH. Measures of *V̇*CO_2_ were corrected for drift using baseline CO_2_ measurements taken at the start and end of each trace and divided by flow rate. We determined the mean *V̇*CO_2_ for the final ten minutes of each trace (Expedata Software, v 1.2.02, Sable Systems).

### Data analysis

We used an information theoretic approach and (generalised) linear mixed models ((G)LMM) for data analysis. For all analyses we created a full model, null model containing only the intercept and random factors, and alternative models containing all subsets of fixed factors whilst retaining the same random factors. We assessed model fit using the Akaike Information Criterion (AIC; Burnham & Anderson, 2002), whereby lower AIC values indicate better fit; where multiple potential best models were within ΔAIC = 2.00 of each other, we selected the simplest model as the final model (Burnham & Anderson, 2002). We then estimated each parameter estimate and its 95% profile likelihood confidence interval (CI) from the final model using the ‘confint()’ function in lme4, and used the CI to infer the statistical significance of each predictor in the final model (Nakagawa & Cuthill, 2007).

#### Learning and memory

To confirm that our task induced learning of the CS+ − US association as expected, we combined data from both experiments and used a GLMM with conditioned response (extension of the proboscis) as the response, with trial number, odour type (CS+), stimulus type (CS+ or CS-) and experiment as covariates. We included individual bee as a random factor and used a binomial error structure (link function = “logit”). We analysed memory separately for each experiment, due to differences in timepoints and control groups between experiments. For each experiment we used a binomial GLM with response to odour as the response, and treatment group, timepoint of memory test, odour type and stimulus as covariates.

#### Sucrose consumption

We first looked at the rate of sucrose and water consumption, by dividing the total volume consumed by the length (in minutes) that consumption was measured for that individual. We adjusted both sucrose and water consumption for evaporation. We used a GLM/LM with rate of sucrose/water consumption as the response, and treatment and individual mass as covariates. For the model of sucrose consumption, we fitted the model using a Gamma distribution (link function = “log”) due to the non-normality of residuals.

We next explored cumulative sucrose consumption. Measures of cumulative volume every ten minutes were temporally autocorrelated, and the autocorrelation function was not improved when auto-regressive functions were fitted. We therefore used only the final cumulative volume of sucrose consumed for bees that reached the 20-hour endpoint to ascertain whether cumulative volume was different between treatments. We built a linear model (LM) with sucrose volume (adjusted for evaporation) as the response, and treatment and individual dry body mass as covariates. We then further split the total volumes of sucrose consumed, adjusted for evaporation, into three discrete time periods (0-1 hour, 1-4 hours, 4-20 hours) and used a second LM to ascertain whether volume consumed (as the response) was predicted by treatment, timepoint, or their interaction, and individual dry body mass. We initially included a random factor for individual bee in this model but removed it as it explained none of the variance and led to singularity errors.

Finally, we used bees in the learning treatment only to determine whether sucrose consumption was predicted by performance in the memory test. We built a GLM with rate of sucrose consumption adjusted for evaporation as the response, and conditioned response (extension of the proboscis to the conditioned stimulus) and timepoint of memory test (24-hours or 72-hours, set as a factor) as predictors. We used a Gamma distribution due to the non-normality of residuals and removed one influential data point based on a Cook’s distance > 1 (Zuur et al., 2009).

#### Metabolic rate

We used the rate of carbon dioxide (*V̇*CO_2_) production as a proxy for metabolic rate. We first checked that there was no difference in *V̇*CO_2_ across experimental days or between later-assigned treatment groups (initial metabolic rate was measured prior to any treatment). We used an LM with mean *V̇*CO_2_ as the response, and experiment day, treatment group and group mass as covariates. We then calculated the percentage change in *V̇*CO_2_ between the baseline *V̇*CO_2_ measurement (pre-treatment) and *V̇*CO_2_ measures at each timepoint (1-, 4-, 24- and 72-hours).

To determine whether *V̇*CO_2_ changed between treatments, we used a LMM with *V̇*CO_2_ percentage change as the response, and treatment, timepoint set as a factor, an interaction between treatment and timepoint, and group mass as covariates. We included a random effect for group, as the same groups were measured across multiple timepoints. We used pre-treatment *V̇*CO_2_ measures (set to 0) as the reference. We repeated this analysis with timepoint as a continuous variable square-root transformed.

Finally, we used bees in the learning treatment only to determine whether initial *V̇*CO_2_ (measured before conditioning trials) predicted performance in a memory test. We built an LM with *V̇*CO_2_ as the response, and conditioned response (extension of the proboscis to the conditioned stimulus) and timepoint of memory test (4-, 24- or 72-hours, set as a factor) as the predictors. Including group as a random factor caused convergence warnings and was removed.

All analyses were carried out in R (v 4.4.1, R Core Team, 2023) using the packages car (Fox & Weisberg, 2019) and dplyr (Wickham et al., 2023) for data manipulation, lme4 (Bates et al., 2015), RVAideMemoire (Herve, 2023), DHARMa (Hartig, 2022), performance (Lüdecke et al., 2021) and MuMIn (Bartoń, 2023) for model building and validation, and ggplot2 (Wickham, 2016), gghalves (Tiedemann, 2022), ggbeeswarm (Clarke, Sherrill-Mix & Dawson, 2023) and patchwork (Pedersen, 2023) for data visualisation.

## Results

### Learning and memory performance

We trained bees to associate a novel odour with a sucrose reward in an associative learning task (n = 82 individual bees in Experiment 1 and n = 115 in Experiment 2). The proportion of correct choices significantly increased across trials (GLMM, trial parameter estimate: 0.35, 95% confidence intervals (CI): 0.30 to 0.41; Fig. 2A; Table S1). When comparing responses of bees to the learnt odour (CS+) versus a second unrewarded odour (CS-), the proportion of bees responding with extension of the proboscis was significantly higher to the rewarded versus unrewarded odour, indicating learning had occurred (GLMM, stimulus parameter estimate: 2.80, 95% CIs: 2.40 to 3.22; Fig. 2A). Learning success varied between odour type, with bees trained to lemon and orange odours as the CS+ making significantly more correct choices compared with aniseed, but there was no difference in response probability between ginger and aniseed odours (GLMM, lemon parameter estimate: 1.29, 95% CIs: 0.42 to 2.17; orange parameter estimate: 1.03, 95% CI: 0.20 to 1.88, ginger parameter estimate: 0.48, 95% CIs: −0.38 to 1.36). There was no difference in the number of correct choices between Experiments 1 and 2 (experiment was not retained in the best model). Thus, overall, bees successfully learnt to form positive associations with a specific odour and discriminated between a learnt rewarding and unrewarding odour.

**Figure 2.**
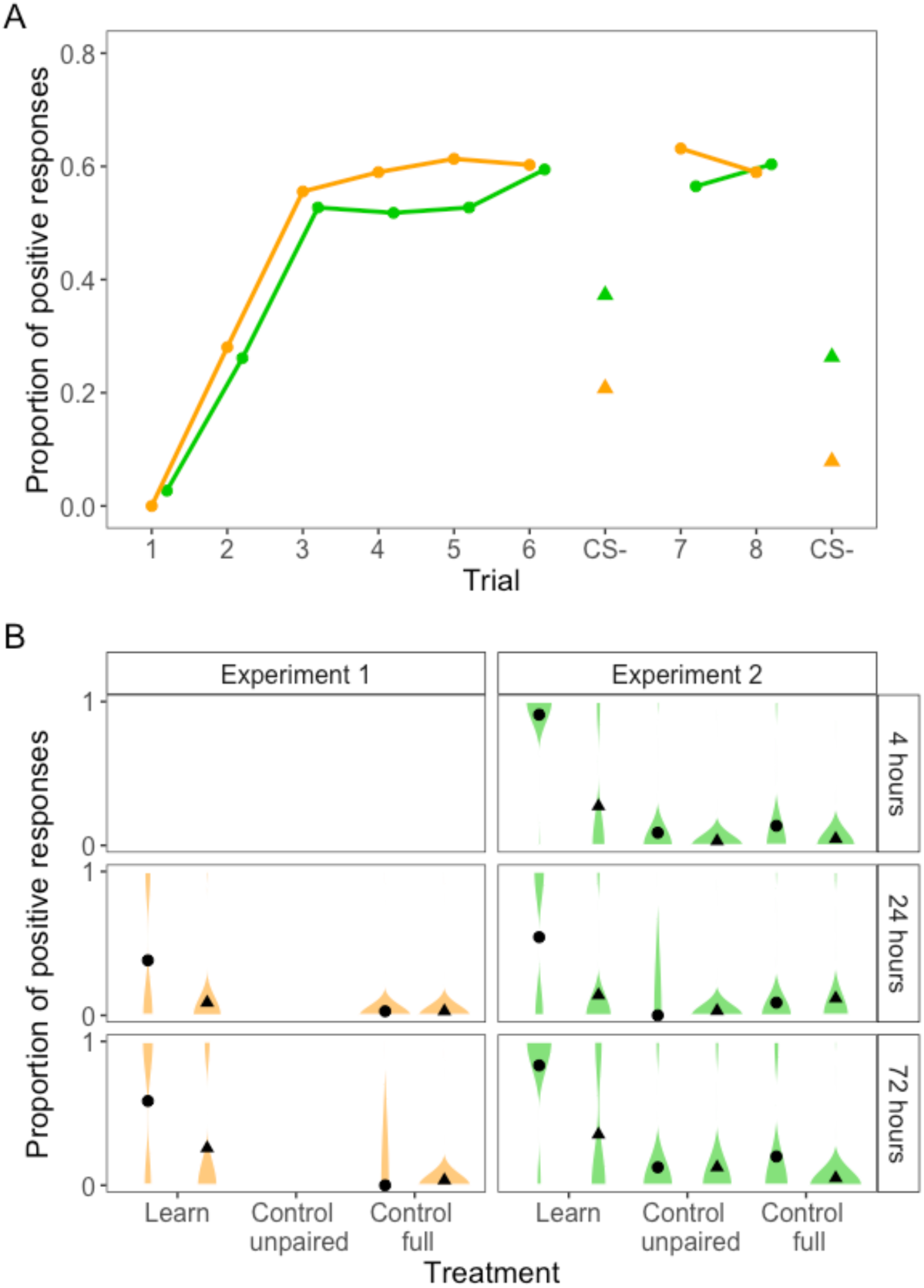
(A) Proportion of positive responses made by honeybees in an olfactory associative conditioning task in two separate experiments (Experiment 1 – orange, Experiment 2 – green). Between trials 6 and 7, and after trial 8, we presented bees with an unrewarded odour (CS-, triangle). (B) Proportion of positive responses during memory tests to a previously rewarded (CS+, circle) or unrewarded (CS-, triangle) odour at 4-, 24- or 72-hours after conditioning across bees in the learning or control treatments. Bees in the unpaired control were exposed to the odour and reward unpaired, so that they could not form a positive association. Bees in the full control were not exposed to the odour but received an equivalent reward volume. Shaded violins show raw data counts. Note that in Experiment 1 we did not test memory at 4-hours, and there was no unpaired control group. The number of bees tested at each timepoint were: Experiment 1, learners at 24h = 41, 72h = 39, control-full at 24h = 38, 72h = 36; Experiment 2, learners at 4h = 24, 24h = 65, 72h = 20, control-unpaired at 4h = 36, 24h = 40, 72h = 25, control-full at 4h = 25, 24h = 45, 72h = 30.

Memory trials were completed at 24- and 72-hours (Experiment 1), and 4-, 24- and 72-hours (Experiment 2) following learning trials. Across both experiments, bees in the learning group had a significantly higher number of positive responses to any odour presentation compared with control bees (GLM, learning treatment parameter estimate and 95% CIs; Experiment 1: 3.09, 2.01 to 4.55; Experiment 2: 2.49, 1.83 to 3.21; Fig. 2B; Table S2, S3) and positive responses were significantly higher for previously rewarded (CS+) versus unrewarded (CS-) odours (GLM, CS+ stimulus parameter estimate and 95% CIs; Experiment 1: 1.33, 0.57 to 2.14; Experiment 2: 1.71, 1.16 to 2.30; Fig. 2B). For Experiment 2, there was no difference in responses between the full control and unpaired control groups (GLM, unpaired treatment parameter estimate: −0.86, 95% CIs: −1.79 to 0.02).

The probability of responding to the correct odour during memory tests differed between timepoints. For Experiment 1, positive responses were slightly, but significantly, higher for memory tests at 72-hours compared with 24-hours (GLM, 72-hour timepoint parameter estimate: 0.80, 95% CIs: 0.06 to 1.57). For Experiment 2, positive responses were significantly lower for tests at 24-hours compared with tests at 4-hours (GLM, 24-hour timepoint parameter estimate: −0.87, 95% CIs: −1.52 to −0.24), but there was no difference in responses between the 4-hour and 72-hour timepoints (GLM, 72-hour timepoint parameter estimate: 0.64, 95% CIs: −0.10 to 1.39). Therefore, across both experiments, memory recall at 24-hours appeared poorer compared with recall at the 4-hour or 72-hour timepoints.

The probability of making a correct choice varied with odour type in memory tests for Experiment 2. There was a significantly lower proportion of correct choices made with ginger (GLM, parameter estimate: −1.53, 95% CIs: −2.45 to −0.69) and orange (GLM, parameter estimate: −1.46, 95% CIs: −2.39 to −0.59) odours compared with aniseed, but no difference in responses between lemon and aniseed odours (GLM, parameter estimate: −0.31, 95% CIs: −0.92 to 0.29). There was no difference in responses to odour types for memory tests in Experiment 1 (odour type was not retained in the best model), likely due to fewer odours being used for learning and memory trials in this experiment.

### Experiment 1: Effects of memory formation on sucrose consumption

Overall, the total volume of sucrose consumed at 20-hours was significantly higher for bees in the learning treatment compared with control bees (Fig. 3A; LM, learning parameter estimate: 11.35, 95% CIs: 4.16 to 18.55, control treatment n = 28 bees, learning treatment n = 30 bees; Table S4). Accordingly, bees in the learning treatment had a non-significant higher rate of total sucrose consumption compared with bees in the control group (mean±SEM learning treatment (n = 75 bees): 0.39±0.16 µL min^-1^, control treatment (n = 67 bees): 0.15±0.04 µL min^-1^; GLM, learning parameter estimate: 0.98, 95% CIs: −0.01 to 1.95; Table S5). When looking at cumulative consumption, the increase in consumption by bees in the learning treatment appeared around the 5-hour timepoint (Fig. 3A). There was no effect of bee body mass on sucrose consumption (body mass was not retained in the best model, in neither the model for sucrose consumption rate, nor total consumption at 20-hours).

**Figure 3.**
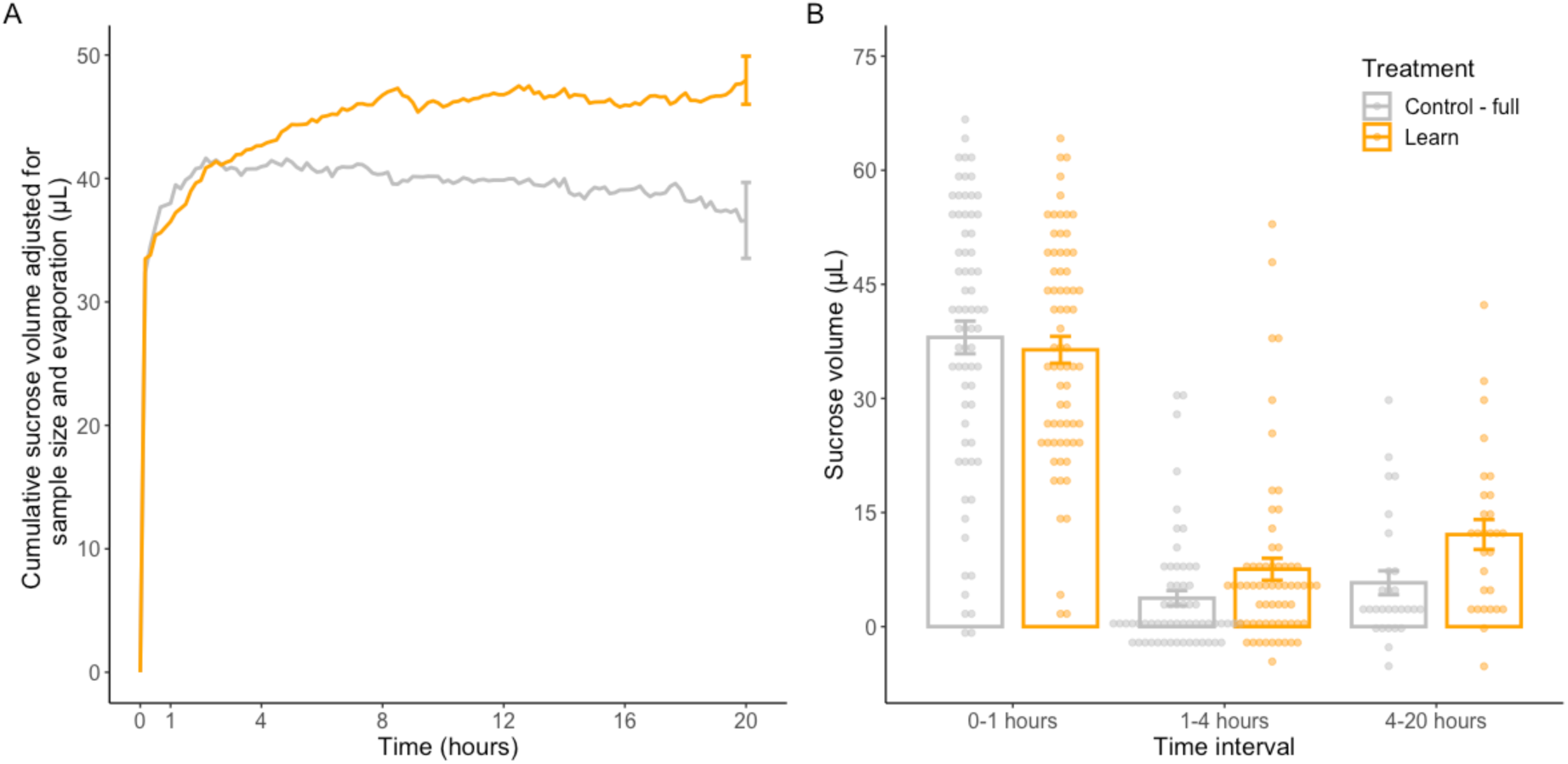
(A) Cumulative sucrose consumption by bees in the learning (orange) or control (grey) treatment groups measured every 10 minutes over a 20-hour period, corrected for evaporation. Lines shows the mean, error bars show the standard error. Note that not all bees persisted for the full 20-hour period (starting sample size, learner treatment = 81, control treatment = 79 bees; end-point sample size, learner treatment = 28, control treatment = 30 bees), which results in a downward trend in mean cumulative volume consumed over time as mean sucrose consumption is corrected for sample size. (B) Total sucrose volume consumed by bees at 0-1-hours, 1-4-hours and 4-20-hours after conditioning trials. Bars show the mean and standard error, points show the raw data. Volumes are adjusted for evaporation, resulting in some negative consumption values.

We grouped sucrose consumption into three discrete timepoints (0-1 hours, 1-4 hours and 4-20 hours). We based these timepoints on previous work in fruit flies showing that consumption increases in the 0-4-hours compared with 4–20-hour intervals (Plaçais et al., 2017). We further included the initial 0– 1-hour time interval, to account for the large volume of sucrose consumed by bees in both groups during this period. For both learner and control bees, consumption was highest in the first hour, and consumption for both treatment groups at the 1-4-hour and 4–20-hour timepoints was significantly lower than at the 0–1-hour interval (LM, 1–4-hour parameter estimate: −31.56, 95% CIs: −34.73 to - 28.39, 4-20-hour parameter estimate: −27.82, 95% CIs: −31.87 to −23.78; Fig. 3B; Table S6). At the 1-4-hour and 4–20-hour timepoints, consumption was higher for bees in the learning treatment, but this was not significant (treatment, or the interaction between treatment and time, was not retained in the best model; Fig. 3B). Consumption was also higher for bees with a larger dry body mass across both treatment groups (LM, mass parameter estimate: 1.72, 95% CIs: 0.27 to 3.16). There was no difference in the rate of water consumption between bees in the learning or control treatment (mean±SEM 0.01±0.00 µL min^-1^; LM, treatment or body mass were not retained in the best model, Table S7). When looking only at bees in the learning treatment, the probability of a positive response in memory tests did not explain rates of sucrose consumption (Fig. S2A; LM, memory score or timepoint were not retained in the best model, null model accepted; Table S8).

### Experiment 2: Effects of memory formation on metabolic rate

We first looked at baseline measures of the rate of carbon dioxide production (*V̇*CO_2_) as a proxy for standard metabolic rate, taken before groups of bees were allocated to treatments. *V̇*CO_2_ did not vary with time (experiment day), later-assigned treatment group, or group body mass (these covariates were not retained in the best model; Table S9).

We next explored percentage change in *V̇*CO_2_, using *V̇*CO_2_ measured before conditioning trials as the baseline (set to 0). Across all groups, there was a significant effect of time on *V̇*CO_2_ percentage change. *V̇*CO_2_ significantly increased at the 1-hour timepoint (Fig. 4; LMM, 1-hour parameter estimate: 7.95, 95% CIs: 1.42 to 14.48; Table S10) and significantly decreased at the 4-hour timepoint (LMM, 4-hour parameter estimate: −16.35, 95% CIs: −23.50 to −9.19). *V̇*CO_2_ percentage change at the 24-hour and 72-hour timepoints did not significantly differ from 0 (LMM, 24-hour parameter estimate: 0.55, 95% CIs: −7.17 to 8.31, 72-hour parameter estimate: −1.79, 95% CIs: −17.58 to 14.06). Due to low survival rates, few groups were measured at the 72-hour timepoint (learner groups n = 3, control-unpaired n = 3, control-full n = 1), but removing this timepoint from the analysis did not change the result (see Supplementary Results; Table S11). Group body mass had no effect on *V̇*CO_2_ percentage change (mass was not retained in the best model).

**Figure 4.**
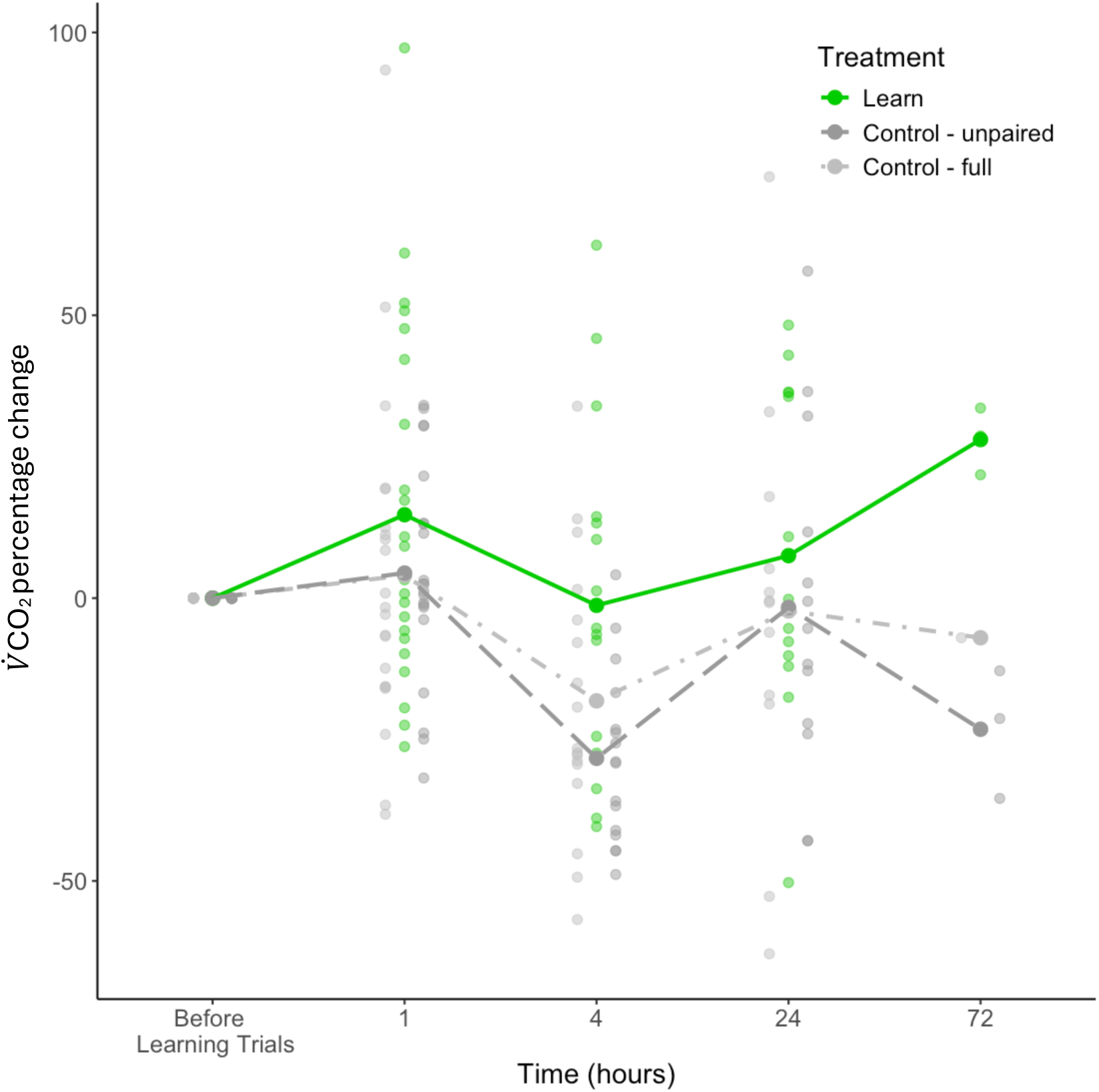
Percentage change of mean metabolic rate (measured as rate of carbon dioxide production, *V̇*CO_2_) of honeybees pooled in groups of five workers, measured before learning trials, and then at 1-hour, 4-hours, 24-hours and 72-hours after learning trials. Groups were either assigned to a learning treatment (olfactory conditioning, green solid line), an unpaired control group in which bees were presented with the odour but unpaired from the sucrose reward (dark grey dashed line) and a full control group in which bees were not presented with the odour but were fed an equivalent volume of sucrose (light grey dashed line). Points show the raw data.

Groups that underwent learning trials had a higher *V̇*CO_2_ percentage change across all timepoints, but this difference was not statistically significant (Fig. 4; treatment, or the interaction between treatment and time, was not retained in the best model). The largest difference appeared at the 4-hour timepoint; there was a decrease in metabolic rate across all treatment groups, but learners decreased by only 1% (n = 16 groups), compared with a −18% and −28% change for the unpaired (n = 16) and full (n = 17) control groups, respectively. For groups in the learning treatment, *V̇*CO_2_ percentage change at the 1-hour and 24-hour timepoints was positive (15% (n = 23) and 8% (n = 14) increase, respectively). Control groups saw a smaller increase at the 1-hour timepoint (4% (n = 21) for both unpaired and full control groups), and a decrease in *V̇*CO_2_ compared to the baseline at the 24-hour timepoint (−2% for both unpaired (n = 13) and full (n = 12) control groups). Thus, overall, changes in metabolic rate followed different trajectories across groups, but this did not result in a significantly higher metabolic rate overall for those in the learning group. The result did not change when timepoint was included as a continuous variable in the model. For this model, only treatment was retained in the best model, but there was no significant effect of treatment on change in metabolic rate (learning parameter estimate: 9.58, 95% CIs: −0.12 to 19.25, unpaired control parameter estimate: −1.98, 95% CIs: −11.82 to 7.93; Table S12; Table S13).

When looking only at bees in the learning treatment, the probability of a positive response to the conditioned stimulus in memory tests did not explain variation in metabolic rate (Fig. S2B, LM, *V̇*CO_2_ was not retained in the best model, null model accepted; Table S14).

## Discussion

We investigated potential energetic costs of long-term memory (LTM) formation in honeybee foragers, measuring sucrose consumption and standard metabolic rate following a spaced associative conditioning protocol. Bees exposed to our conditioning protocol successfully formed LTM that could be recalled after 72-hours, and those in the learning treatment consumed a higher volume of sucrose, but not water, in the 20-hours following conditioning compared with control bees. However, there was no significant increase in metabolic rate following LTM formation. Overall, our findings suggest that formation of LTM in bees drives increased dietary sucrose intake requirements, but these are not substantial enough to significantly influence metabolic rate at the level of the whole organism.

In honeybees, LTM can be divided into two physiologically distinct phases, which are translation (early-LTM) and transcription (late-LTM) dependent, and last for approximately 1-2 days, and 3-4 days, respectively (Menzel, 2012). In both experiments, recall of the learnt odour was higher at the 72-hour versus the 24-hour timepoint, suggesting performance was stronger for transcription-dependent late-LTM within our task/cohort. The timings of transcription during memory formation are likely to reflect increases in energy requirements. Two phases of transcription are associated with the formation of LTM across taxa (Alberini, 2009), and have been identified in honeybees. An initial phase lasting approximately 40 minutes occurs during learning trials, followed by a second phase that lasts up to nine hours after conditioning (Lefer et al., 2013). The increase in sucrose consumption that we observed around five hours after conditioning trials may, therefore, correspond with the second phase of transcription during late-LTM formation, and we assume that synthesis of proteins and transcription factors involved in synaptic reorganisation may occur during this time (Alberini & Kandel, 2015).

Our results echo those of Plaçais et al. (2017), who showed that fruit flies doubled their sucrose consumption within the first four hours following spaced olfactory conditioning compared with flies in a control group. However, our increase in 1.3× the volume of sucrose consumed by bees in the learning versus control treatments at the 20-hour timepoint is smaller and likely driven by differences in the concentration of solution provided (our sucrose solution had an 8× higher concentration compared with the sucrose solution provided to fruit flies because bees do not typically respond to low concentrations in learning assays).

Metabolic rate responds to short-term changes in energetic demands. For example, temporary increases in metabolic rate have previously been documented during periods of digestion (Secor, 2009), immune response (Ardia et al., 2012), and flight (Kammer & Heinrich, 1978). We therefore expected that metabolic rate may respond to increases in energy demands during LTM formation. However, we found no effect of spaced olfactory conditioning on rate of CO_2_ production (as a proxy for standard metabolic rate) in groups of bees. Our results suggest that increases in energy demands at the cellular level may not translate to detectable changes in metabolic rate at the level of the whole organism. Given that energy demands of LTM formation are localised to specific neurons in the mushroom bodies (Plaçais et al., 2017), and that metabolic rate is sensitive to all other body processes, large changes in metabolic rate following memory formation may be unlikely following formation of a single memory, as occurred in our paradigm. However, it is also possible that our protocol was not sensitive to very small trends in metabolic rate. For example, to increase the sensitivity of our equipment, we measured metabolic rate in groups of five bees that had all undergone the same treatment, which were caught at hive entrances as they returned from foraging bouts, thus we could not standardise for inter-individual differences in foraging behaviour or experience, and any potential effects on metabolic rate. It is, therefore, possible that inter-individual variation in metabolic rate within a group masked relatively smaller differences between treatments. Repeating the study with equipment that allows for metabolic rate measurements for individual bees could clarify whether the observed non-significant increase in metabolic rate across learning groups is driven by individual variation or investment in memory and warrants further investigation.

Across all treatment groups, we observed that metabolic rate decreased when measured at the 4-hour timepoint following conditioning protocols. Patterns in metabolic rate following digestion have been identified in most animals, including insects (Secor, 2009). The 4-hour timepoint in our experiment corresponds with the longest time since feeding for honeybees (bees were fed each morning and during conditioning), so it is unlikely that bees were actively digesting at this timepoint, resulting in a relatively lower metabolic rate. Overall, metabolic rate was influenced more by energetic state (i.e., recent sucrose consumption) over any potential changes in energy demands of appetitive LTM formation. Future studies could explore whether formation of aversive (cf. appetitive) LTM differently affects metabolic rate, removing the use of sugar solution as a reward during learning trials. Furthermore, it would be interesting to contextualise the effects of starvation on the relationship between LTM formation and metabolic rate. Glucose is the main source of energy for the brain (Sokoloff, 1999), but periods of starvation may induce alternative energy-synthesis pathways. For example, under starvation, glial cells synthesise lipids to form ketone bodies as an alternative energy source (Silva et al., 2022), but the potential effects on metabolic rate remain unexplored.

Metabolic rate has been implicated as a potential driver of intra-specific variation in behaviour (Biro & Stamps, 2010). For example, individuals with a higher resting metabolic rate may have to increase foraging behaviours to support their relatively higher energy demands, compared to conspecifics with lower resting metabolic rates (Mathot, Dingemanse & Nakagawa, 2019). Accordingly, positive correlations have been identified between resting metabolic rate and behaviours that may aid net energy-gain during foraging, such as boldness and dominance (Brown et al., 2003; Finstad et al., 2007; McCarthy, 2001). For bees, appetitive LTM is relevant during foraging as individuals form positive associations between floral traits and rewards (Menzel, 1993). However, we found no relationship between group metabolic rate taken before learning trials (as a measure of baseline metabolic rate) and individual memory score, suggesting that variation in metabolic rate did not explain variation in appetitive memory abilities. The relationship between metabolic rate and aversive LTM may be different, as aversive memories are less strongly associated with behaviours that result in net energy-gain (Mathot, Dingemanse & Nakagawa, 2019). Repeating the experiment using an aversive conditioning protocol could provide useful insights into differences in energetic costs between appetitive and aversive conditioning protocols. A potential limitation of our experimental protocol is that individuals learnt a single association, but learning in the wild is likely to involve different sensory modalities and more than one association (Menzel, 1993). A more complex learning paradigm may have revealed relationships between memory and metabolic rate. Nonetheless, our protocol using a single olfactory association is relevant to honeybees, which show high degrees of specialisation on a single floral species during a foraging bout (Chittka, Thomson & Waser, 1999; Grüter & Ratnieks, 2011).

### Conclusions

Energy availability has been predicted as a constraint on the evolution of neural systems (Niven & Laughlin, 2008), therefore energetic costs may be important to explain variation in cognitive abilities, both intra- and inter-specifically (Boogert et al., 2018; Thornton & Lukas, 2012). Our results suggest that costs at the cellular level may not have detrimental impacts to the whole organism when energy can be compensated for through increased dietary intake. Induced costs of synaptic transmission and potential synaptic plasticity during LTM formation may, therefore, not have a significant effect on the overall energy budget of the brain, above baseline synaptic activity (Karbowski, 2019). However, the impact of a cost associated with LTM formation may become important when energy is limited. Further work is needed to determine how the expression of learning and memory traits might change in response to changing energetic states of an individual, and contexts in which individuals continue to invest in cognitive abilities when under energetic stress.

## Acknowledgements

We thank Alex Clarke and students from the Department of Electronic Engineering, Royal Holloway University of London for building and programming the Arduino controller, Sean Gibson for making capillary tubes for sucrose consumption measurements, Keith McMahon for apiary maintenance, and Abbie Reade for sharing information about the CAFE assay. We are grateful to Adam Watrobski for support with equipment transport and Catherine Charnock from Manchester Cathedral for allowing us access to their bees during a pilot experiment. C.M.W. is funded by the ICL-RHUL BBSRC DTP (BB/M011178/1).

## Data availability

All data and code are available at: doi.org/10.6084/m9.figshare.28123643.

