## Supplementary Information for "Long-term memory formation impacts dietary energy intake but not metabolic rate in honeybees"

### Supplementary Results

For the main metabolic rate model that asked whether treatment affects $\dot{V}CO_2$ percentage change, we re-ran the model with the 72-hour timepoint removed, due to low samples sizes (learner group n=3, unpaired control group n=3, full control group n=1 bees). The results did not change, and $\dot{V}CO_2$ significantly increased at the 1-hour timepoint (LMM, parameter estimate: 7.95, 95% CIs: 1.37 to 14.52) and decreased at the 4-hour timepoint (LMM, parameter estimate: -16.46, 95% CIs: -23.65 to -9.25) compared with the pre-treatment baseline. There was no difference between $\dot{V}CO_2$ at the 24-hour timepoint compared with the pre-treatment baseline (LMM, parameter estimate: 0.58, 95% CIs: -7.19 to 8.38). Treatment, the interaction between treatment and timepoint, and group mass were not retained in the best model (Table S11).

Removing the 72-hour timepoint from the model where timepoint was treated as a continuous variable also did not alter the result, and the null model was accepted (Table S13).

Supplementary Figures

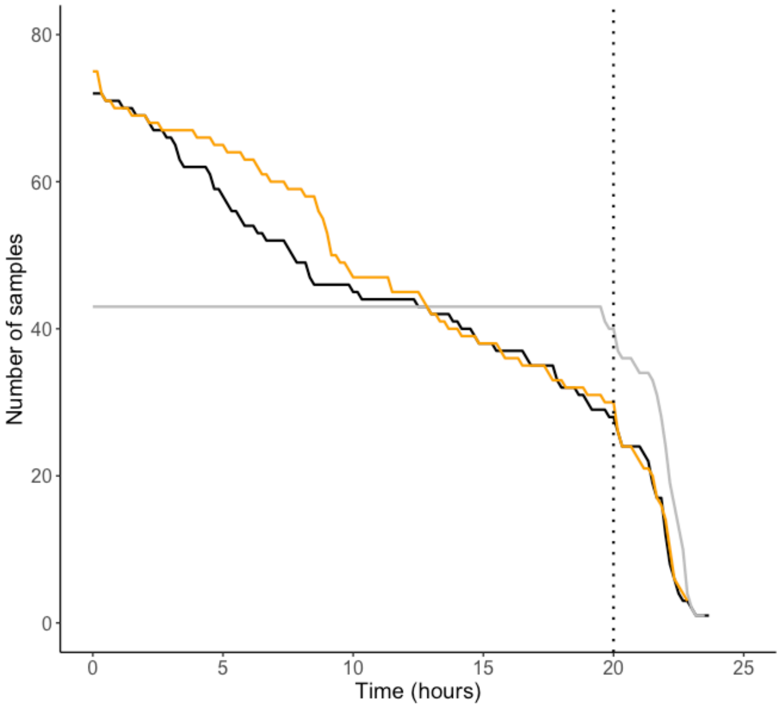


**Figure S1.** Number of samples (videos of sucrose consumption) at each timepoint (10-minute intervals) for bees in the learning treatment (orange), control (black) and evaporation control (grey). Videos were stopped when bees had drunk all available sucrose or water, were unable to access the sucrose or water, or when there was a leak. We used 20-hours as the cut-off for all bees (vertical black dotted line).

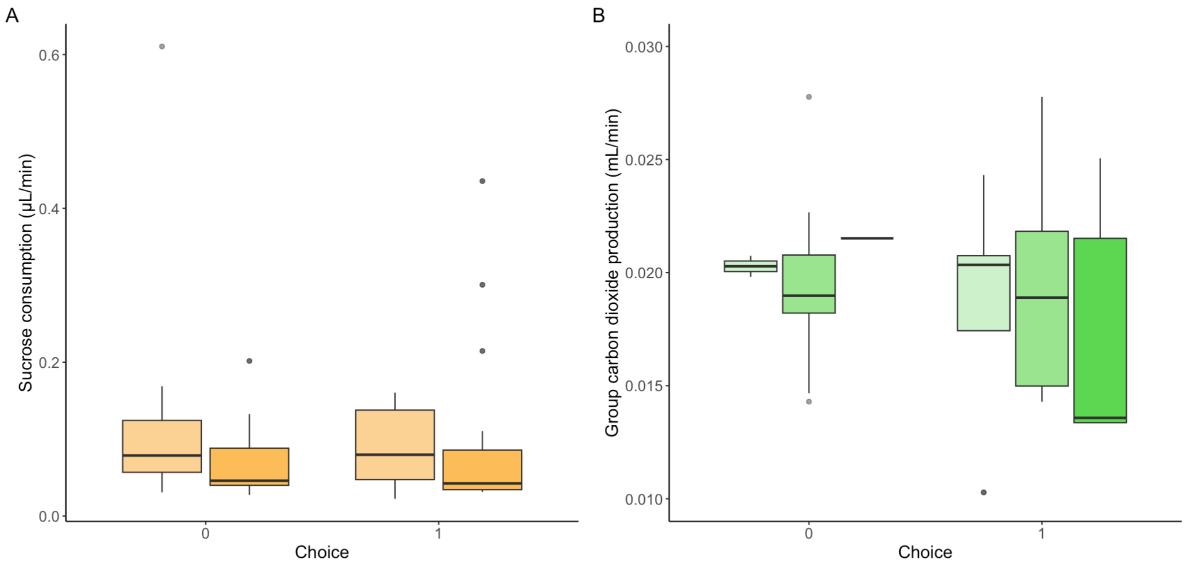


**Figure S2.** Outcomes of memory assays across both experiments (0 = no response to conditioned stimulus, 1 = positive response to conditioned stimulus). (A) Memory was tested at 24-hours (light orange) and 72-hours (dark orange), and performance on the memory task did not predict rate of sucrose consumption in the 20-hours following conditioning. (B) Memory was tested at 4-hours (lightest green), 24-hours (light green), and 72-hours (dark green), and performance on the memory task did not predict rate of carbon dioxide production (as a proxy for standard metabolic rate).

Supplementary Tables

For all statistical models, the model in bold shows the final model.

**Table S1.** Candidate model set for a generalised linear mixed model investigating olfactory learning in bees, with probability of extension of the proboscis (conditioned response, 0/1) as the response, and trial number (trials 1-10), odour type (aniseed/ginger/lemon/orange essential oils), experiment year (Experiment 1 = 2021, Experiment 2 = 2023), and stimulus type (positive (rewarded) conditioned stimulus or negative (unrewarded) conditioned stimulus) as covariates. The model was fitted using a binomial error structure with individual bee as a random effect.

| **Covariates:** trial number + odour + stimulus + year<br>**Random effect:** individual bee | | | |
|---|---|---|---|
| Model | DF | AIC | ΔAIC |
| **Odour + Trial + Stimulus** | **7** | **2003.4** | **0.00** |
| Trial + Odour + Year + Stimulus | 8 | 2005.4 | 2.00 |
| Trial + Stimulus | 4 | 2008.1 | 4.70 |
| Trial + Year + Stimulus | 5 | 2010.1 | 6.70 |
| Odour + Stimulus | 6 | 2193.7 | 190.34 |
| Odour + Year + Stimulus | 7 | 2195.7 | 192.33 |
| Stimulus | 3 | 2198.8 | 195.41 |
| Year + Stimulus | 4 | 2200.8 | 197.40 |
| Trial + Odour | 6 | 2230.3 | 226.93 |
| Trial + Odour + Year | 7 | 2232.3 | 228.93 |
| Trial | 3 | 2234.9 | 231.52 |
| Trial + Year | 4 | 2236.9 | 233.52 |
| Odour | 5 | 2272.5 | 269.09 |
| Odour + Year | 6 | 2274.5 | 271.09 |
| Null | 2 | 2277.3 | 273.93 |
| Year | 3 | 2279.3 | 275.93 |

**Table S2.** Candidate model set for a generalised linear model investigating olfactory memory in bees (Experiment 1), with probability of extension of the proboscis (conditioned response, 0/1) as the response, and treatment (learner group or full control group), memory timepoint (24-hours and 72-hours after conditioning), learnt odour (positively reinforced odour during learning trials, orange/ginger), and stimulus (positive (rewarded) conditioned stimulus or negative (unrewarded) conditioned stimulus) as covariates. The model was fitted using a binomial error structure.

| **Covariates:** treatment + timepoint + learnt odour + stimulus | | | |
|---|---|---|---|
| Model | DF | AIC | ∆AIC |
| **Stimulus + Timepoint + Treatment** | **4** | **180.7** | **0.00** |
| Treatment + Timepoint + Learnt odour + Stimulus | 5 | 182.5 | 1.81 |
| Treatment + Stimulus | 3 | 183.2 | 2.49 |
| Treatment + Learnt odour + Stimulus | 4 | 185.0 | 4.23 |
| Treatment + Timepoint | 3 | 190.9 | 10.23 |
| Treatment + Timepoint + Learnt odour | 4 | 192.8 | 12.06 |
| Treatment | 2 | 192.8 | 12.12 |
| Treatment + Learnt odour | 3 | 194.6 | 13.89 |
| Timepoint + Stimulus | 3 | 225.3 | 44.61 |
| Timepoint + Learnt odour + Stimulus | 4 | 227.1 | 46.41 |
| Stimulus | 2 | 227.8 | 47.05 |
| Learnt odour + Stimulus | 3 | 229.6 | 48.85 |
| Timepoint | 2 | 233.8 | 53.09 |
| Timepoint + Learnt odour | 3 | 235.6 | 54.91 |
| Null | 1 | 235.8 | 55.07 |
| Learnt odour | 2 | 237.6 | 56.89 |

**Table S3.** Candidate model set for a generalised linear model investigating olfactory memory in bees (Experiment 2), with probability of extension of the proboscis (conditioned response, 0/1) as the response, and treatment (learner group, unpaired control group or full control group), memory timepoint (4-hours, 24-hours and 72-hours after conditioning), learnt odour (positively reinforced odour during learning trials, aniseed/orange/ginger/lemon), and stimulus (positive (rewarded) conditioned stimulus or negative (unrewarded) conditioned stimulus) as covariates. The model was fitted using a binomial error structure.

| **Covariates:** treatment + timepoint + learnt odour + stimulus | | | |
|---|---|---|---|
| Model | DF | AIC | ΔAIC |
| **Treatment + Timepoint + Learnt odour + Stimulus** | **9** | **382.9** | **0.00** |
| Stimulus + Timepoint + Treatment | 6 | 397.2 | 14.38 |
| Treatment + Learnt Odour + Stimulus | 7 | 399.2 | 16.34 |
| Treatment + Stimulus | 4 | 407.6 | 24.72 |
| Treatment + Timepoint + Learnt odour | 8 | 421.6 | 38.76 |
| Treatment + Timepoint | 5 | 434.2 | 51.29 |
| Treatment + Learnt odour | 6 | 436.1 | 53.23 |
| Treatment | 3 | 443.2 | 60.33 |
| Time + Learnt odour + Stimulus | 7 | 498.9 | 116.02 |
| Learnt odour + Stimulus | 5 | 502.5 | 119.61 |
| Stimulus | 2 | 507.0 | 124.16 |
| Timepoint + Stimulus | 4 | 507.5 | 124.64 |
| Timepoint + Learnt odour | 6 | 528.1 | 145.19 |
| Learnt odour | 4 | 531.3 | 148.46 |
| Null | 1 | 535.4 | 152.55 |
| Timepoint | 3 | 536.0 | 153.17 |

**Table S4.** Candidate model set for a linear model investigating the effect of treatment group (learner or control group) and individual dry mass on the total volume of sucrose consumed (µL) over a 20-hour period following olfactory conditioning.

| **Covariates:** treatment + dry mass | | | |
|---|---|---|---|
| Model | DF | AIC | ΔAIC |
| **Treatment** | **3** | **471.9** | **0.00** |
| Treatment + Dry mass | 4 | 472.2 | 0.33 |
| Null | 2 | 479.4 | 7.52 |
| Dry mass | 3 | 479.5 | 7.61 |

**Table S5.** Candidate model set for a generalised linear model investigating the effect of treatment group (learner or control group) and individual dry mass on the rate of sucrose consumption (µL/min). The model was fitted using a Gamma error structure to account for overdispersion.

| **Covariates:** treatment + dry mass | | | |
|---|---|---|---|
| Model | DF | AIC | ∆AIC |
| Treatment + Dry mass | 4 | -146.9 | 0.00 |
| **Treatment** | **3** | **-146.7** | **0.22** |
| Dry mass | 3 | -133.8 | 13.08 |
| Null | 2 | - 130.8 | 16.10 |

**Table S6.** Candidate model set for a linear model investigating the effect of treatment group (learner or control group), timepoint (0-1, 1-4, or 4-20 hours), a treatment:timepoint interaction, and individual dry mass on the total volume of sucrose consumed (µL) during each discrete timepoint. Dry mass was normalised using the scale function in R.

| **Covariates:** treatment + timepoint + treatement:timepoint + dry mass | | | |
|---|---|---|---|
| Model | DF | AIC | ∆AIC |
| Treatment + Timepoint + Treatment:Timepoint + Dry mass | 8 | 2548.5 | 0.00 |
| **Timepoint + Dry mass** | **5** | **2548.9** | **0.33** |
| Treatment + Timepoint + Dry mass | 6 | 2549.7 | 1.15 |
| Treatment + Timepoint + Treatment:Timepoint | 7 | 2551.8 | 3.22 |
| Timepoint | 4 | 2552.4 | 3.83 |
| Treatment + Timepoint | 5 | 2552.7 | 4.12 |
| Dry mass | 3 | 2819.1 | 270.58 |
| Null | 2 | 2820.3 | 271.74 |
| Treatment + Dry mass | 4 | 2820.9 | 272.34 |
| Treatment | 3 | 2821.8 | 273.30 |

**Table S7.** Candidate model set for a linear model investigating the effect of treatment group (learner or control group) and individual dry mass on the rate of water consumption (µL/min).

| **Covariates:** treatment + dry mass | | | |
|---|---|---|---|
| Model | DF | AIC | ΔAIC |
| **Null** | **2** | **-539.1** | **0.00** |
| Dry mass | 3 | -537.4 | 1.73 |
| Treatment | 4 | -537.2 | 1.89 |
| Treatment + Dry mass | 4 | -535.6 | 3.57 |

**Table S8.** Candidate model set for a generalised linear model investigating the effect of memory score on the rate of sucrose consumption (µL/min) using only bees that underwent olfactory conditioning. The model included memory score (probability of extension of the proboscis to the conditioned odour, 0/1) and memory timepoint set as a factor (24-hours or 72-hours) as covariates. The model was fitted using a Gamma error structure to account for overdispersion.

| **Covariates:** memory score + memory timepoint + memory score:timepoint | | | |
|---|---|---|---|
| Model | DF | AIC | ΔAIC |
| **Null** | **2** | **-154.7** | **0.00** |
| Memory timepoint | 3 | -153.6 | 1.10 |
| Memory score | 3 | -152.7 | 1.96 |
| Memory score + Memory timepoint | 4 | -91.4 | 63.31 |

**Table S9.** Candidate model set for a linear model investigating the effects of experiment day, treatment (learner, unpaired control, or full control groups) and group dry mass on mean group rate of $CO_2$ production measured before learning and memory trials (as a proxy for baseline metabolic rate). N=5 honeybees per group.

| **Covariates:** experiment day + treatment + group mass | | | |
|---|---|---|---|
| Model | DF | AIC | ΔAIC |
| Experiment day | 3 | -512.9 | 0.00 |
| Experiment day + Treatment | 5 | -512.0 | 0.87 |
| **Null** | **2** | **-511.6** | **1.23** |
| Treatment | 4 | -511.4 | 1.45 |
| Experiment day + Group mass | 4 | -511.0 | 1.86 |
| Experiment day + Treatment + Group mass | 6 | -510.4 | 2.49 |
| Treatment + Group mass | 5 | -510.2 | 2.67 |
| Group mass | 3 | -510.1 | 2.81 |

**Table S10.** Candidate model set for a linear mixed model investigating the effects of treatment (learner, unpaired control, or full control groups), timepoint (before conditioning, and 1-hour, 4-hours, 24-hours or 72-hours after conditioning), an interaction between treatment and timepoint, and group dry mass, on percentage change in mean group rate of $CO_2$ production. Baseline rate of $CO_2$ production (measured before conditioning trials) was set to 0 and used to compare all other timepoints. Individual group was included as a random effect. Timepoint was set as a factor. N=5 honeybees per group.

**Covariates:** treatment + timepoint + treatment:timepoint + group dry mass
**Random effect:** group

| Model | DF | AIC | ΔAIC |
|---|---|---|---|
| Treatment + Timepoint | 9 | 2037.1 | 0.00 |
| **Timepoint** | **7** | **2038.8** | **1.67** |
| Treatment + Timepoint + Group mass | 10 | 2038.9 | 1.80 |
| Timepoint + Group mass | 8 | 2040.7 | 3.61 |
| Treatment + Timepoint + Treatment:Timepoint | 17 | 2040.9 | 3.80 |
| Treatment + Timepoint + Treatment:Timepoint + Group mass | 18 | 2042.7 | 5.66 |
| Treatment | 5 | 2069.0 | 31.92 |
| Treatment + Group mass | 6 | 2070.8 | 33.69 |
| Null | 3 | 2071.2 | 34.10 |
| Group mass | 4 | 2073.1 | 36.03 |

**Table S11.** Candidate model set for a linear mixed model investigating the effects of treatment (learner, unpaired control, or full control groups), timepoint (before conditioning, and 1-hour, 4-hours, 24-hours after conditioning), an interaction between treatment and timepoint, and group dry mass, on percentage change in mean group rate of $CO_2$ production. The model was based on Table S10 with the 72-hour timepoint removed due to a low sample size. Baseline rate of $CO_2$ production (measured before conditioning trials) was set to 0 and used to compare all other timepoints. Individual group was included as a random effect. N=5 honeybees per group.

| **Covariates:** treatment + timepoint + treatment:timepoint + group dry mass<br>**Random effect:** group | | | |
|---|---|---|---|
| Model | DF | AIC | ΔAIC |
| Treatment + Timepoint | 8 | 1974.9 | 0.00 |
| **Timepoint** | **6** | **1975.6** | **0.74** |
| Treatment + Timepoint + Group mass | 9 | 1976.7 | 1.89 |
| Timepoint + Group mass | 7 | 1977.6 | 2.72 |
| Treatment + Timepoint + Treatment:Timepoint | 14 | 1978.7 | 3.89 |
| Treatment + Timepoint + Treatment:Timepoint + Group mass | 15 | 1980.6 | 5.77 |
| Treatment | 5 | 2008.4 | 33.58 |
| Null | 3 | 2009.6 | 34.79 |
| Treatment + Group mass | 6 | 2010.3 | 35.44 |
| Group mass | 4 | 2011.6 | 36.76 |

**Table S12.** Candidate model set for a linear mixed model investigating the effects of treatment (learner, unpaired control, or full control groups), timepoint (before conditioning, and 1-hour, 4-hours, 24-hours or 72-hours after conditioning), an interaction between treatment and timepoint, and group dry mass, on percentage change in mean group rate of $CO_2$ production. Baseline rate of $CO_2$ production (measured before conditioning trials) was set to 0 and used to compare all other timepoints. Individual group was included as a random effect. Timepoint was included as a continuous variable with a square-root transformation. N=5 honeybees per group.

| **Covariates:** treatment + timepoint + treatment:timepoint + group dry mass<br>**Random effect:** group | | | |
|---|---|---|---|
| Model | DF | AIC | ΔAIC |
| **Treatment** | **5** | **2069.0** | **0.00** |
| Treatment + Timepoint | 6 | 2070.3 | 1.30 |
| Treatment + Timepoint + Treatment:Timepoint | 8 | 2070.3 | 1.34 |
| Treatment + Group mass | 6 | 2070.8 | 1.77 |
| Null | 3 | 2071.2 | 2.18 |
| Treatment + Timepoint + Group mass | 7 | 2072.1 | 3.05 |
| Treatment + Timepoint + Treatment:Timepoint + Group mass | 9 | 2072.2 | 3.16 |
| Timepoint | 4 | 2072.5 | 3.47 |
| Group mass | 4 | 2073.1 | 4.12 |
| Timepoint + Group mass | 5 | 2074.4 | 5.39 |

**Table S13.** Candidate model set for a linear mixed model investigating the effects of treatment (learner, unpaired control, or full control groups), timepoint (before conditioning, and 1-hour, 4-hours, 24-hours after conditioning), an interaction between treatment and timepoint, and group dry mass, on percentage change in mean group rate of $CO_2$ production. The model was based on Table S12 with the 72-hour timepoint removed due to a low sample size. Baseline rate of $CO_2$ production (measured before conditioning trials) was set to 0 and used to compare all other timepoints. Individual group was included as a random effect. Timepoint was included as a continuous variable with a square-root transformation. N=5 honeybees per group.

| **Covariates:** treatment + timepoint + treatment:timepoint + group dry mass<br>**Random effect:** group | | | |
|---|---|---|---|
| Model | DF | AIC | ∆AIC |
| Treatment | 5 | 2008.4 | 0.00 |
| Treatment + Timepoint | 6 | 2009.5 | 1.06 |
| **Null** | **3** | **2009.6** | **1.21** |
| Treatment + Group mass | 6 | 2010.3 | 1.86 |
| Timepoint | 4 | 2010.7 | 2.24 |
| Treatment + Timepoint + Group mass | 7 | 2011.3 | 2.88 |
| Group mass | 4 | 2011.6 | 3.19 |
| Treatment + Timepoint + Treatment:Timepoint | 8 | 2012.1 | 3.64 |
| Timepoint + Group mass | 5 | 2012.6 | 4.20 |
| Treatment + Timepoint + Treatment:Timepoint + Group mass | 9 | 2013.9 | 5.48 |

**Table S14.** Candidate model set for a linear model investigating the effect of memory score on mean group $CO_2$ production measured before conditioning trials (as a proxy for baseline metabolic rate). The model included memory score (probability of extension of the proboscis to the conditioned odour, 0/1), memory timepoint (4-hours, 24-hours or 72-hours), and an interaction between memory score and timepoint as covariates.

| **Covariates:** memory score + timepoint + memory score:timepoint | | | |
|---|---|---|---|
| Model | DF | AIC | ∆AIC |
| **Null** | **2** | **-785.6** | **0.00** |
| Memory score | 3 | -785.1 | 0.48 |
| Timepoint | 4 | -783.1 | 2.45 |
| Memory score + Timepoint | 5 | -782.2 | 3.39 |
